# Slow chromatin dynamics enhances promoter accessibility to transcriptional condensates

**DOI:** 10.1101/2021.02.16.431394

**Authors:** Tetsuya Yamamoto, Takahiro Sakaue, Helmut Schiessel

## Abstract

Enhancers are DNA sequences at a long genomic distance from target genes. Recent experiments suggest that enhancers are anchored to the surfaces of condensates of transcription machinery and that the loop extrusion process enhances the transcription level of their target genes. Here we theoretically study the polymer dynamics driven by the loop extrusion of the linker DNA between an enhancer and the promoter of its target gene to calculate the contact probability of the promoter to the transcription machinery in the condensate. Our theory predicts that when the loop extrusion process is active, the contact probability increases with increasing linker DNA length. This finding reflects the fact that the relaxation time, with which the promoter stays in proximity to the surface of the transcriptional condensate, increases as the length of the linker DNA increases. This contrasts the equilibrium case for which the contact probability between the promoter and the transcription machineries is smaller for longer linker DNA lengths.

## Introduction

Enhancers are short regulatory DNA sequences that activate the transcription of target genes. Enhancers are located a long genomic distance (in the order of 10k - 1Mbps) away from target gene promoters and the linker genome thus has to form loops to drive the interactions between enhancers and gene promoters. Mediator complexes that bind to enhancers recruit RNA polymerase II (Pol II) to promoters^1^ and promote the assembly of preinitiation complexes. ^2,3^

Hi-C experiments have shown that genomes of eukaryotic cells are composed of topologically associated domains (TADs),^4–6^ which are indeed chromatin loops of the order of 10k - 1Mbps.^7^ The influence of chromatin loops on the promoter-enhancer interactions was studied by molecular dynamics simulations that assume chromatin as a (semi)flexible polymer with static loops at equilibrium.^8,9^ In contrast to these assumptions, recent theories predict that chromatin loops are produced by the loop extrusion process, with which cohesin acts as a molecular motor that uni-directionally transports chromatin to increase the size of loops until it collides with boundary elements, such as CTCF.^10,11^ Mediator complexes may also act as boundary elements.^11,12^ The loop extrusion theory captures the features of the contact frequency map, which is determined by Hi-C experiments.^10,11^ In earlier single molecule experiments, the motor activity of yeast and human cohesin was not detected ^13–15^ and alternative mechanisms of loop extrusion process were proposed.^16,17^ However, the motor activity and the loop extrusion process were directly observed from human and *Xenopus* cohesin in recent single molecule experiments.^18,19^ It is of interest to theoretically predict the roles played by the dynamics of chromatin looping due to the loop extrusion process in the promoter-enhancer interactions and the regulation of gene expression.

Rao and coworkers used auxin-inducible degron technique, which degrades cohesin in response to a dose of auxin, to eliminate chromatin loops.^20^ They showed that eliminating chromatin loops did not change the transcription level of most genes (at a time point 6 hours after cohesin degradation), but significantly decreases the transcription level of the target genes of superenhancers, which are genomic regions with high density of enhancers. ^21^ Mediator complexes, transcription factors and Pol II form condensates (which are called transcriptional condensates) due to the phase separation driven by the multivalent interactions between the intrinsically disordered domains of these proteins. ^22–26^ Recent microscopic experiments showed that superenhancers colocalize with transcriptional condensates. ^22,23^ Since the linker DNA between the promoters and enhancers is excluded from the transcriptional condensates,^25,26^ this implies that the superenhancers are localized at their surfaces. ^22,23^ These experiments suggest that the loop extrusion of chromatin at the surfaces of transcriptional condensates plays a key role in enhancing the transcription level of the target genes of superenhancers.

We have therefore theoretically analyzed the dynamics of chromatin, which is extruded by cohesin, at the surface of a transcriptional condensate. ^27,28^ First, by using a bead-spring model we predicted that the mean square end-to-end distance of a chromatin region decreases with a constant rate in the bulk solution, whereas it does not decrease until the tension generated by cohesin, extruding the chain from the grafted end, reaches the free end.^27^ Second, we used Onsager’s variational approach to predict that the loop extrusion process increases the local concentration of chromatin at the surface of a transcriptional condensate and the lateral pressure generated by the excluded volume interactions between chromatin units decreases the surface tension of the transcriptional condensate. ^28^ Here we take into account the chromatin dynamics driven by the loop extrusion process in an extension of the Langmuir’s theory of surface adsorption to predict the accessibility of gene promoters to the transcription machineries in a transcriptional condensate. Large transcriptional condensates are stable for the experimental time scale (with which mouse embryonic stem cells are differentiated), whereas the lifetime of small transcriptional condensates is in the order of 10 s.^24^ For simplicity, we limit our discussion to stable transcripition condensates. Our theory predicts that for cases in which the loop extrusion process is effective, the contact probability increases with increasing the length of linker chromatin. This contrasts with the case in which loop extrusion is inhibited, where the contact probability decreases with increasing the length of linker chromatin between the gene promoter and the enhancer. The increased contact probability in the presence of loop extrusion reflects the fact that the relaxation time, with which the gene promoters stay in proximity to the surface of the transcriptional condensate, increases with increasing the length of the linker chromatin. ^29,30^ The longer relaxation time enhances the probability that the gene promoter rebinds to the surface of the transcriptional condensate. The contact probability of gene promoters to transcriptional machineries is proportional to the length of transcription bursts and may be experimentally accessible.

## Model

### Chromatin section anchored at surface of transcriptional condensate

We consider a linker chromatin between a promoter and an enhancer, which is part of a super-enhancer associated to the surface of a transcriptional condensate, see fig. 1a. We treat a stable transcriptional condensate.^24^ The promoter and enhancer have affinities to the transcriptional condensate because of the stochastic binding of transcription factors, whereas the linker chromatin between the promoter and the enhancer is expelled from the condensate.^25,26^ The linker chromatin is composed of *N*_0_ chain units and the promoter is composed of *N_s_* units, where the length of each unit is *b* for both the linker and the promoter. For simplicity, we treat cases in which the number of units in the promoter is smaller than the number of units in the linker chromatin, *N_s_* ≪ *N*_0_. We also neglect the unbinding of enhancers from the condensate.

**Figure 1:**
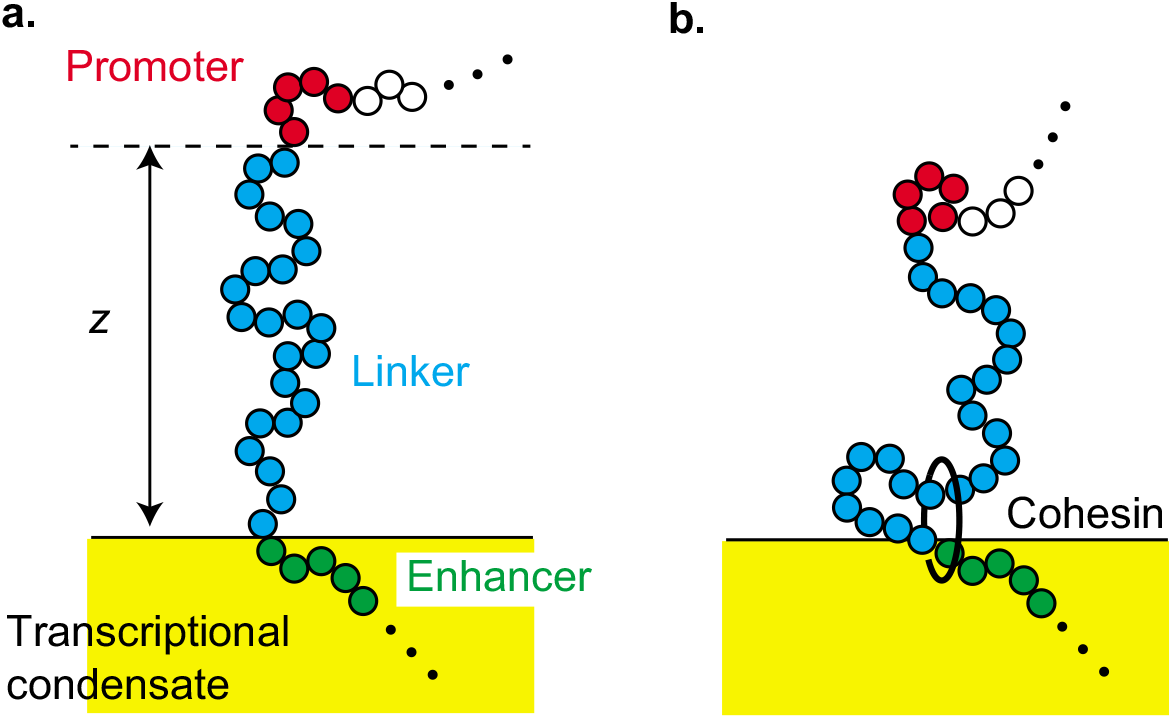
a. Model of a linker chromatin with *N*_0_ units between an enhancer (green) and the promoter (red) of its target gene at the surface of a transcriptional condensate. The enhancer is a part of the super-enhancer and is localized at the surface of a transcriptional condensate. The linker chromatin (blue) between the enhancer and the promoter is expelled from the condensate, b. Cohesin is loaded from the enhancer and translocates chromatin units from the arm region (which has not been extruded) to the loop region (which has been extruded) with a constant rate 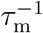. *z* is the distance between the position of the promoter and the surface of the condensate.

In equilibrium, the distribution of the position z of the promoter above the surface of the condensate has the form^31–35^

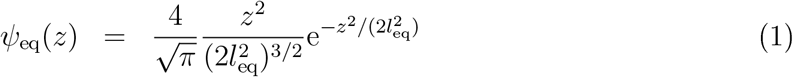

where the mean square end-to-end vector 3*l*^2^ of the linker chromatin is given by

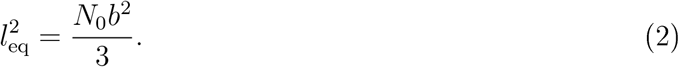

For simplicity, we analyze only the *z*-position of the promoter, i.e. the position normal to the surface. The factor *z*^2^ in eq. (1) results from the repulsive interactions between the linker chromatin and the transcriptional condensate and from the fact that the linker is part of a longer chromatin chain. The derivation of eq. (1) is shown in the supplementary material (and see also refs.^31–35^).

Following refs.^32–34^ we treat the linker chromatin between the promoter and the enhancer as a dumbbell in an effective potential

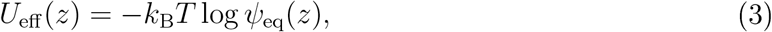

where *k_B_* is the Boltzmann constant and *T* is the absolute temperature. For cases in which the loop extrusion does not influence the polymer dynamics, the probability distribution function *ψ*(*z,t*) of the position *z* of the promoter at time *t* is derived by the Smoluchowski equation

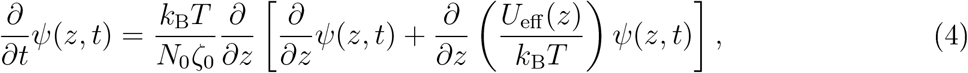

where ζ_0_ is the friction constant of each chromatin unit. The first term of eq. (4) represents the thermal fluctuations (diffusion) of the linker chromatin. The second term of eq. (4) represents the fact that the linker chromatin plays a role in the (entropic) spring of stiffness 3*k*_B_T/(*N*_0_*b*^2^) and the contribution of the effective potential −2*k*_B_*T* log *z* due to the repulsive interactions between the linker chromatin and the surface. The solution of eq. (4) can be derived by using the eigen function expansion, see sec. S2 in the Supplementary File.

### Chromatin dynamics during loop extrusion and relaxation

Cohesin is preferentially loaded at the site, at which NIPBL-MAU2 is localized.^13–15^ Indeed, experiments suggest that the loop extrusion process is driven by the complex of cohesin and NIPBL-MAU2.^18^ Recent experiments showed that TADs are recovered relatively fast at superenhancers, implying that there are active loading sites at super-enhancers. ^20^ The loop extrusion may become asymmetric when the cohesin loading site is at the proximity to the elements that stop the loop extrusion,^36^ such as mediators in the transcriptional condensate.^11,12^ Motivated by the latter experiments, we treat cases in which cohesin is loaded from the enhancer end of the linker chromatin with constant average loading time *τ*_On_. The loop extrusion by cohesin divides the linker chromatin into the loop and arm regions where only the former region has already been extruded by cohesin, see fig. 1b. Cohesin translocates chromatin units from the arm region to the loop region with a constant rate 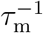.

For simplicity, we assume that a new cohesin starts the loop extrusion process before the old one is unloaded from the chromatin at the end of the TAD but that there is not more than one cohesin in the linker chromatin between the enhancer and the promoter. We set the time at which the promoter is extruded by a cohesin to *t* = 0 *ψ*(*z*, 0) = *δ*(*z*), see fig. 2**a**. The linker chromatin is now included in a loop and the number of chromatin units increases with time *t*

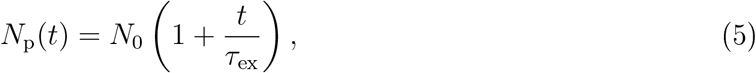

where *τ*_ex_ (= *N*_0_*τ_m_*) is the time scale with which a chromatin section of length *N*_0_ is extruded.

**Figure 2:**
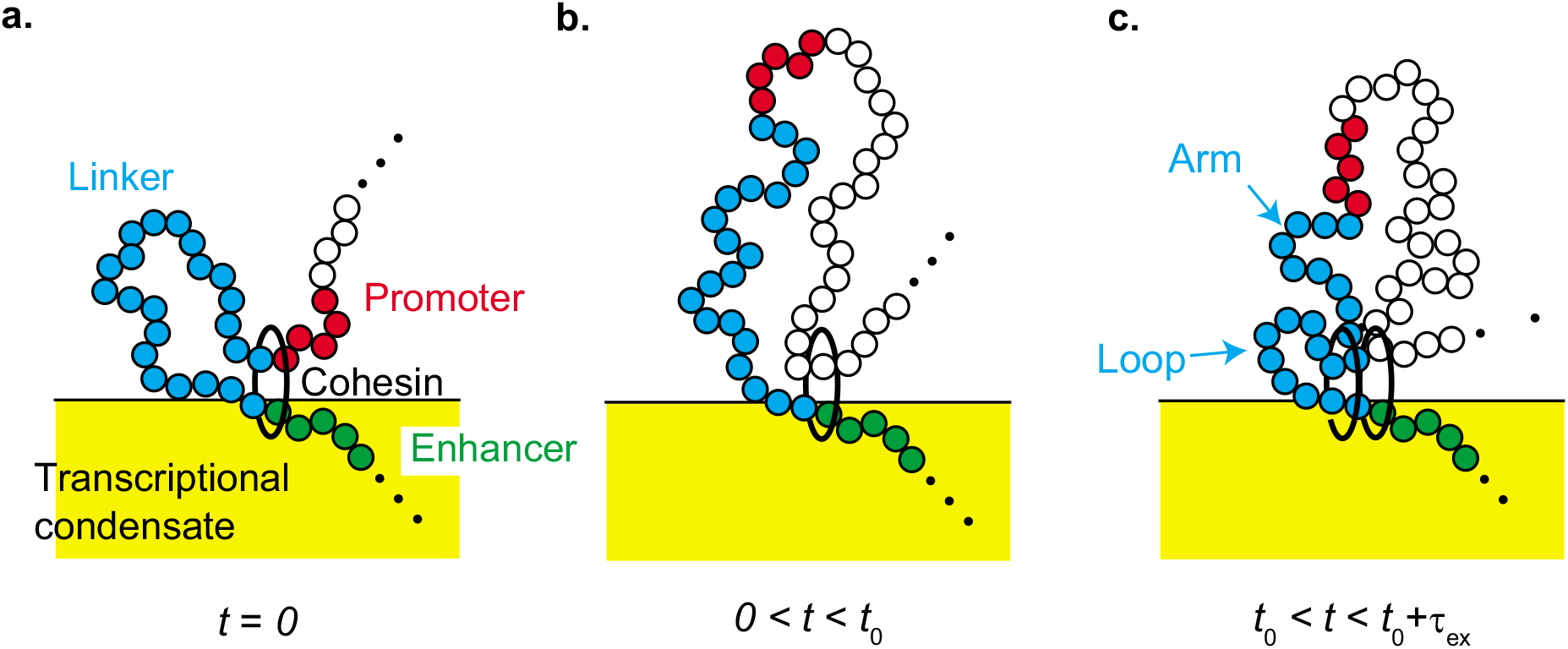
We set *t* = 0 to the time when the gene promoter is translocated to a loop by a cohesin (**a**). The linker chromatin between the promoter and the enhancer relaxes to the (local) equilibrium conformation in the relaxation process, 0 < *t* < *t*_0_ (**b**). A new cohesin is loaded from the enhancer and translocates chromatin units from the arm region (which has not been extruded by the new cohesin) to the loop region (which has been extruded by the cohesin) with a constant rate 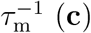 (**c**).

The linker chromatin is relaxed to the (local) equilibrium in the growing loop during 0 < *t* < *t*_0_ (which we call relaxation process). A new cohesin starts the loop extrusion process at *t* = *t*_0_ (*t′* = 0) and the promoter is eventually extruded by the new cohesin at *t* = *t*_0_ + *τ*_ex_ (*t′* = *τ*_ex_) (which we call loop extrusion process). In the following, we assume *t*_0_ = *τ*_on_ to highlight the roles played by the dynamics of linker chromatin, although the loading of cohesin is stochastic and treating it as the Poisson process may be more precise. The number N(t) of chromatin units in the arm region is given by *N*_0_ during the relaxation process and by

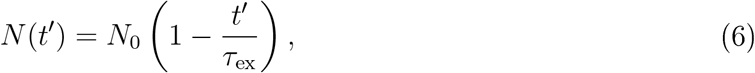

during the loop extrusion process, see fig. 2. The probability distribution function *ψ_loc_*(*z,t*) in the local equilibrium has the form

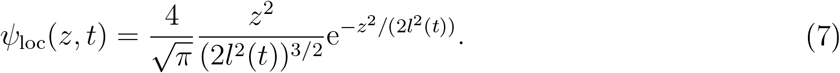

The mean square end-to-end distance 3*l*^2^(*t*) of the linker chromatin at the local equilibrium has the form

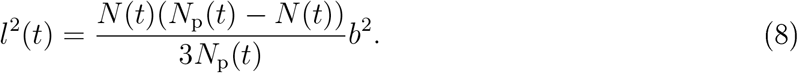

Eq. (7) leads to the effective potential

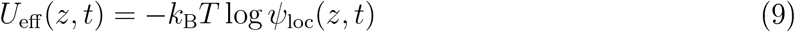

in the relaxation and loop extrusion processes.

During the relaxation process, the time evolution of the probability distribution function *ψ*(*z, t*) of the position *z* of the promoter has the form

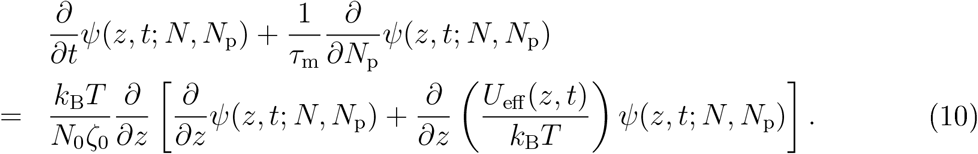

Eq. (10) takes into account the growth of the loop that includes the linker chromatin (see fig. 2**b**) in an extension of eq. (4), see the second term on the left side of eq. (10). During the loop extrusion process, the time evolution of the probability distribution function *ψ*(*z,t*) of the position *z* of the promoter has the form

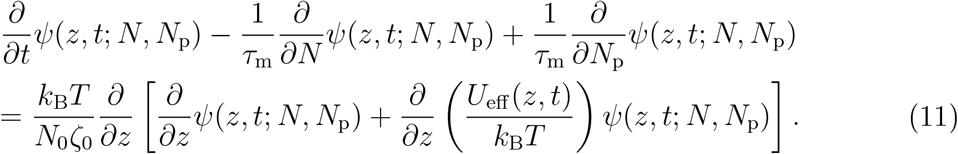

Eq. (11) takes into account the extrusion of chromatin units in the arm region to the loop region (see fig. 2c) in an extension of eq. (10), see the third term on the left side of eq. (11).

### Stochastic binding dynamics of promoter to condensate

The fraction *σ*(*t*) of promoters that bind to the surface of the transcriptional condensate has the form

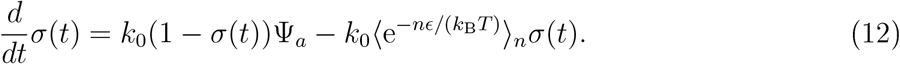

The first term is the binding rate of the promoter to the condensate and the second term is the unbinding rate of the promoter from the condensate. *k*_0_ is the rate constant that accounts for this process. We assume that the promoter binds to the surface of the condensate with a constant rate *k*_0_ when it is located at the reaction zone 0 < *z* < *a* due to the finite size of the promoter, see fig. 3**a**. **Ψ**_*a*_ is the probability with which the promoter is located at the reaction zone

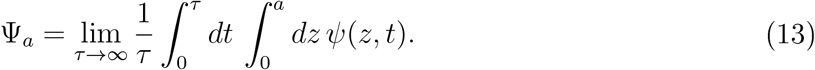

**Figure 3:**
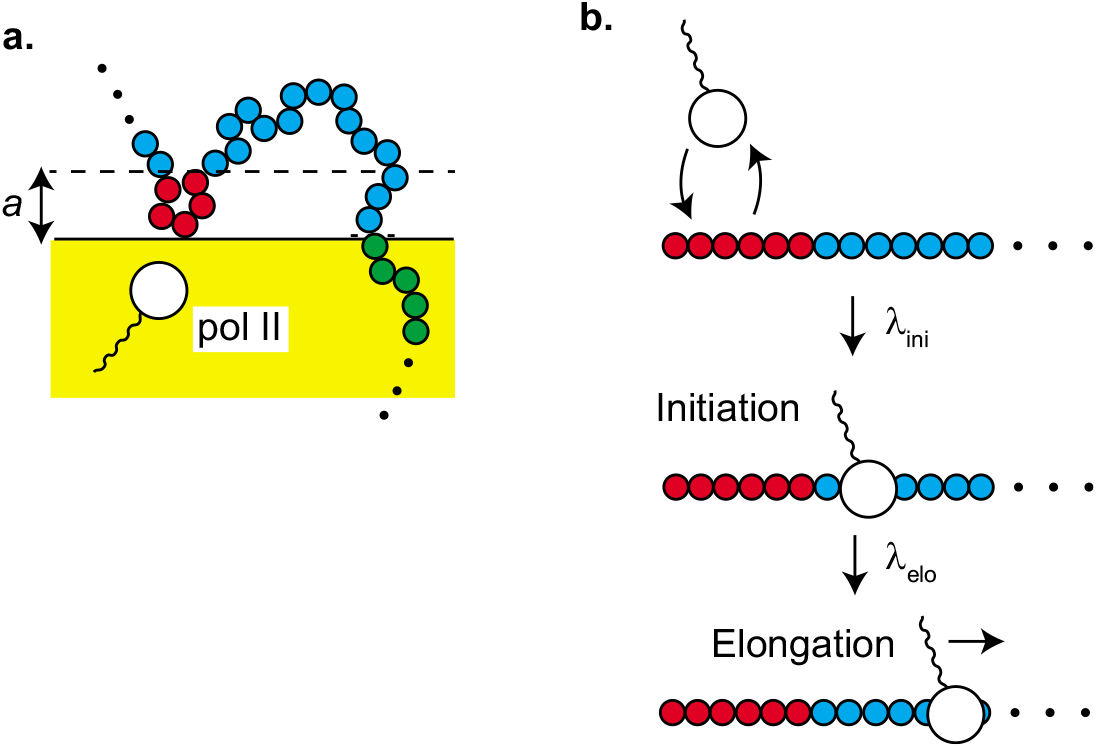
Stochastic binding-unbinding dynamics of promoters at the surface of the transcriptional condensate (**a**). The promoter at the reaction zone 0 < *z* < *a* binds to the surface of the condensate with a constant rate, where *a* is the size of the promoter. RNA polymerase II (Pol II) in the condensate binds to and unbinds from the promoter (**b**). The equilibrium constant *K*_ini_ accounts for the binding-unbinding dynamics. The bound Pol II starts transcription and shows the promoter proximal pause (the initiation state). The rate constant λ_ini_ accounts for the transition to this state. The promoter is tethered to the surface of the condensate by Pol II in the initiation state. Pol II escapes with the rate constant λ_elo_ from the promoter to start elongation.

Experiments by Nozaki and coworkers suggest that Pol II molecules in the initiation state tether the promoter, to which these molecules bind, to the condensate. ^37^ The Boltzmann factor 〈*e*^−*nϵ*/(*k*_B_*T*)^〉_*n*_ accounts for the tethering of the promoter to the condensate by Pol II. *ϵ* is the binding energy between the condensate and Pol II bound to the promoter. 〈〉_*n*_ is the average with respect to the number *n* of Pol II bound to the promoter (in the initiation state, see fig. 3**b** and the discussion below). Eq. (12) therefore takes into account the chromatin dynamics and the tethering of the promoter by Pol II in the initiation state in an extension of the Langmuir’s theory of the dynamics of surface adsorption. ^38^ With eq. (12), we assume that the binding of the gene promoter to the transcriptional condensate is rate limited, motivated by the fact that among the genes that are at the proximity to a condensate and move together with the condensate, only 20 % of them colocalize with the condensate,^24^ see also the discussion.

We simplify a transcription model used by Stasevich and coworkers ^40^ to derive the form of the factor 〈*e*^−*nϵ*/(*k*_B_*T*)^〉_*n*_. Pol II shows stochastic binding and unbinding dynamics to the promoter, see fig. 3**b**. When Pol II bound to the promoter changes its conformation and assembles the preinitiation complex, the enzyme starts transcription and stops ~ 100 nucleotides downstream of the transcription starting site. ^39^ The pausing state of Pol II is called the initiation state. Pol II then starts elongation when its carbon terminal domain is phospholylated, ^39^ The time evolution equation for the probability *P_ini_*(*t*) that the promoter is occupied by Pol II in the initiation state has the form

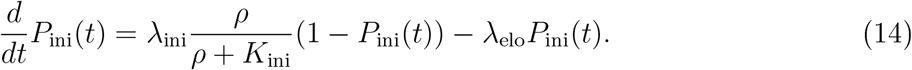

The first term of eq. (14) is the rate with which Pol II enters the initiation state and the second term is the rate with which Pol II enters elongation state, *ρ* is the concentration of Pol II in the transcriptional condensate and *K*_ini_ is the equilibrium constant with respect to the stochastic binding and unbinding dynamics of Pol II to the promoter. λ_ini_ denotes the rate with which Pol II bound to the promoter becomes the initiation state. λ_elo_ is the rate with which Pol II in the initiation state enters the elongation state. For simplicity, we treat cases in which not more than one Pol II can occupy a promoter. We also assume that the difference of Pol II concentration between the interior and exterior of the transcriptional condensate is very large and neglect the transcription by Pol II in the exterior of the transcriptional condensate. We use the probability *P*_ini_(*t*) in the steady state to derive the factor 〈*e*^−*nϵ*/(*k*_B_*T*)^〉_*n*_, assuming that the transcription dynamics is faster than the binding-unbinding dynamics of the promoter to the condensate.

## Results

### Relaxation time of linker chromatin increases as length of linker chromatin increases

To understand the chromatin dynamics during the loop extrusion process and during the relaxation process, we first analyze the distribution function *ψ*(*z,t*) of the position of the promoter. When the promoter is extruded by a cohesin, the promoter is located at the surface of the condensate *ψ*(*z*, 0) = *δ*(*z*). The linker chromatin between the promoter and the enhancer is now included in one loop. The linker chromatin shows the relaxation dynamics towards the equilibrium conformation, while the number of units in the chromatin loop increases with time due to the loop extrusion. As a result the promoter diffuses away from the surface with time due to the thermal fluctuation of the linker chromatin, see fig. 4a. The distribution function eventually becomes that of the equilibrium, if the linker chromatin is completely relaxed before new cohesin is loaded to the linker chromatin, see fig. 4**b**.

**Figure 4:**
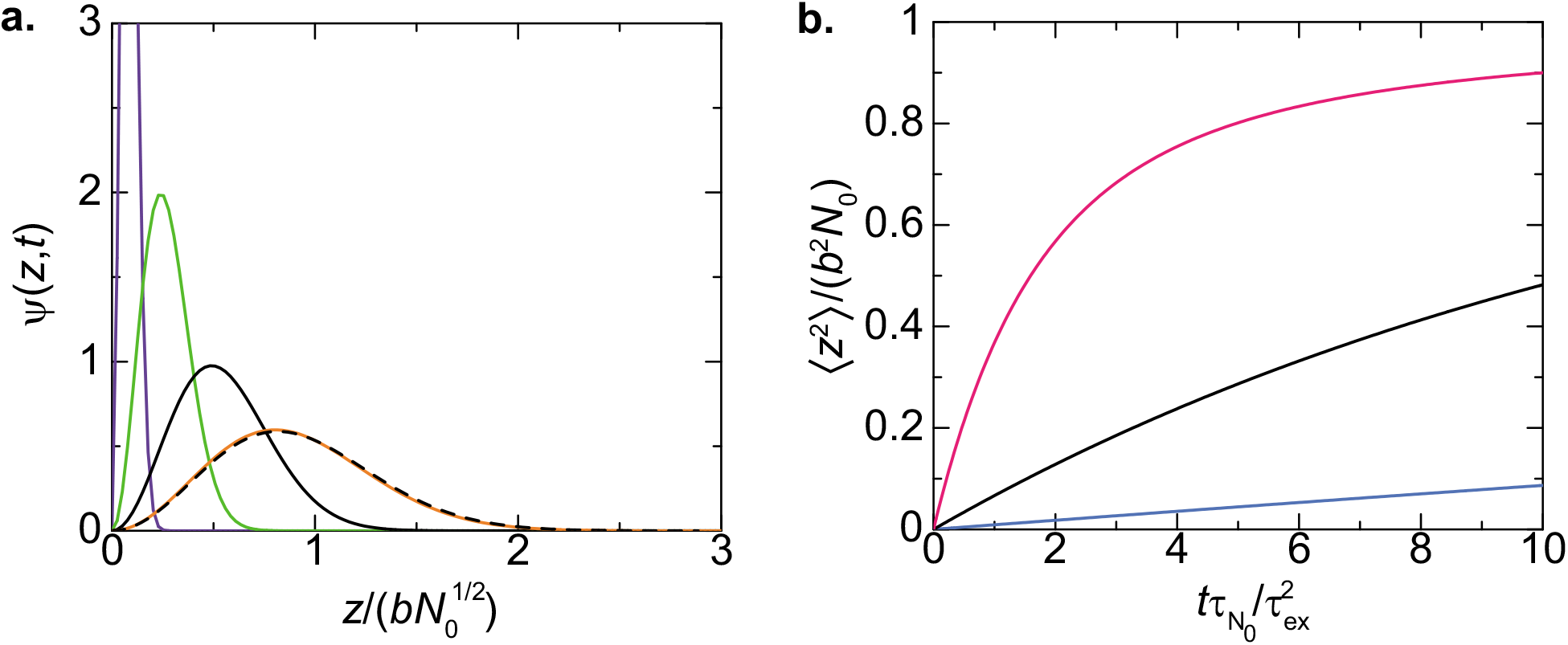
**a** The distribution function *ψ*(*z,t*) of the promoter in the relaxation process is shown as a function of the position *z* of the promoter for 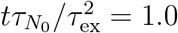 (purple), 10.0 (light green), 50.0 (black), and 500.0 (orange). We used *τ*_ex_/*τ*_*N*_0__ = 0.1. The broken curve is the distribution function at equilibrium, **b.** The mean square distance between the promoter and the surface of the condensate is shown as a function of rescaled time 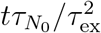 for *τ*_ex_/*τ*_*N*_0__ = 0.1 (cyan), 0.3 (black), and 1.0 (magenta), where the ratio *τ*_ex_/*τ*_*N*_0__ is proportional to the inverse of the number *N*_0_ of units in the linker chromatin. The rescaling factor 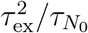 does not depend on the number *N*_0_ of units in the linker chromatin.

The distribution function of the promoter has the form

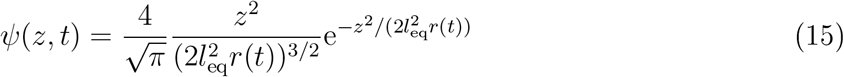

for both the relaxation process and the loop extrusion process, where 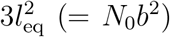 is the mean square distance between the promoter and the enhancer in the equilibrium distribution. The relaxation factor *r*(*t*) is proportional to the mean square distance between the promoter and the enhancer, 〈*z^2^*〉 = *b*^2^*N*_0_*r*(*t*), and its form depends on the process, see eq. (S47) in the Supplementary File for the form of *r*(*t*) in the relaxation process. The relaxation becomes slower as the number *N*_0_ of units in the linker chromatin increases, see fig. 4**b**. The relaxation time has the form

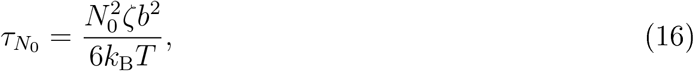

which increases with increasing the number *N*_0_ of units in the linker chromatin. The fact that the relaxation time *τ*_*N*o_ is proportional to the square of the number *N*_0_ of units reflects the Rouse dynamics of the linker chromatin.^29^

A new cohesin is loaded at the enhancer end of the linker chromatin and a new loop extrusion process starts at time *t*_0_. The new cohesin translocates the chromatin units in the arm region to the loop region and thus the promoter is dragged towards the surface of the condensate, see fig. 5. The distribution function of the promoter has the form of eq. (15), but the form of the relaxation factor *r*(*t*) is different from the relaxation process, see eq. (S56) in the Supplementary File. For cases in which the ratio *τ*_ex_/*τ*_*N*_0__ of time scales is very large, the promoter approaches the surface with almost constant rate, see the orange line in fig. 5**b**. In contrast, for small values of the ratio *τ*_ex_/*τ*_*N*_0__, the position of the promoter is hardly affected by the loop extrusion process for a finite period of time before the mean square distance between the promoter and the surface decreases steeply, see the cyan line in fig. 5. This results from the fact that the tension generated by the loop extrusion process diffuses along the linker chromatin and it takes a finite time until the tension drags the promoter towards the surface. This feature agrees well with our previous theory based on a bead-spring model^27^ and Onsager’s variational principle.^28^

**Figure 5:**
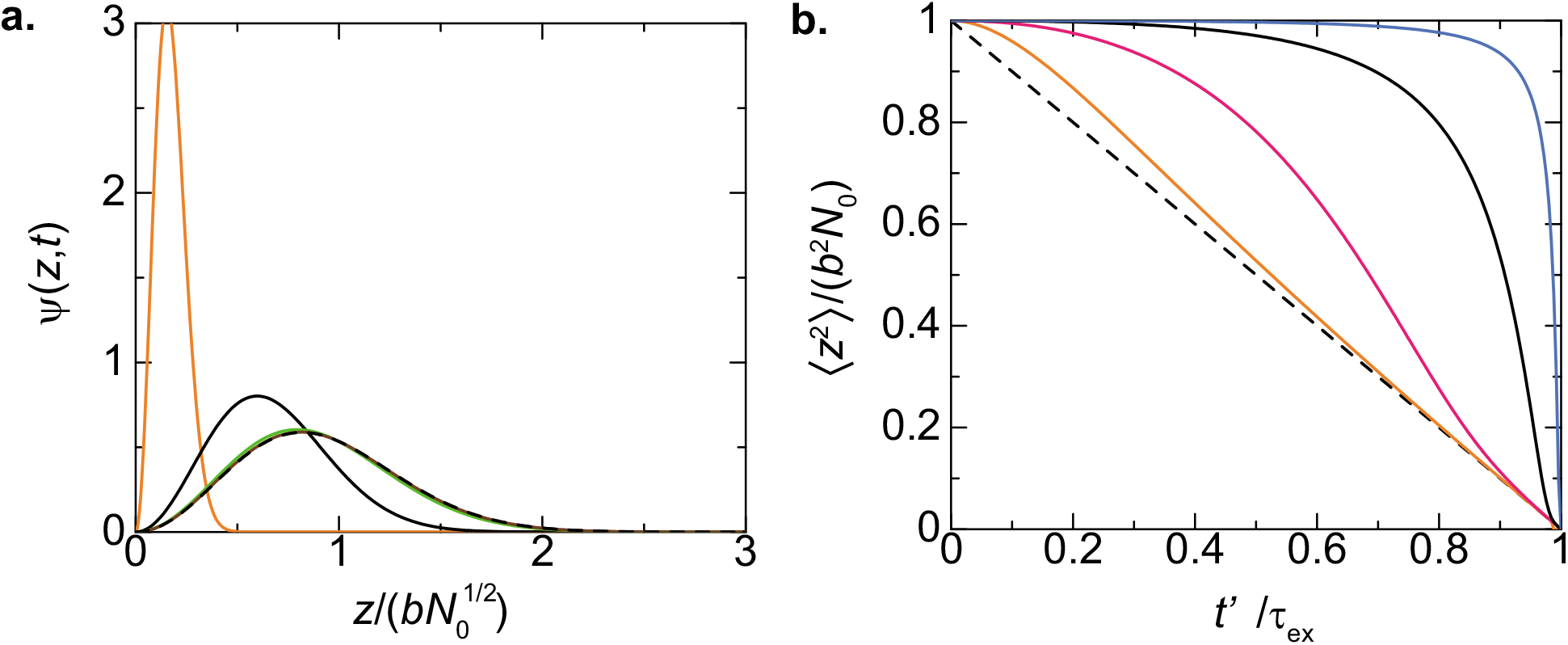
**a.** The distribution function *ψ*(*z, t*) is shown as a function of the position 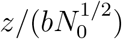 of the promoter for *t′*/*τ*_ex_ = 0.0 (black broken curve), 0.3 (brown), 0.6 (light green), 0.9 (black), and 0.98 (orange), where *t′* (= *t* − *t*_0_) is the time elapsed since the loop extrusion process starts. We used *τ*_ex_/*τ*_*N*_0__ = 0.1. **b.** The mean square distance between the promoter and the surface is shown as a function of time *t′*/*τ*_ex_ for *τ*_ex_/*τ*_*N*_0__ = 0.01 (cyan), 0.1 (black), 1.0 (magenta), and 10.0 (orange). The linker chromatin is completely relaxed (*t*_0_ → ∞) when the loop extrusion starts for both **a** and **b.**

For cases in which the time period *t*_0_ of the relaxation process is larger than the relaxation time *τ*_*N*_0__ of the linker chromatin, the linker chromatin is relaxed to the equilibrium conformation before the loop extrusion process starts, see the magenta line in fig. 6. In contrast, for cases in which the time period *t*_0_ is shorter than the relaxation time *τ*_*N*_0__, the loop extrusion process starts before the linker chromatin is completely relaxed to the equilibrium conformation, see the cyan, light green, and black lines in fig. 6. The promoter thus stays in the proximity of the surface due to the extrusion of the linker chromatin with a constant rate.

**Figure 6:**
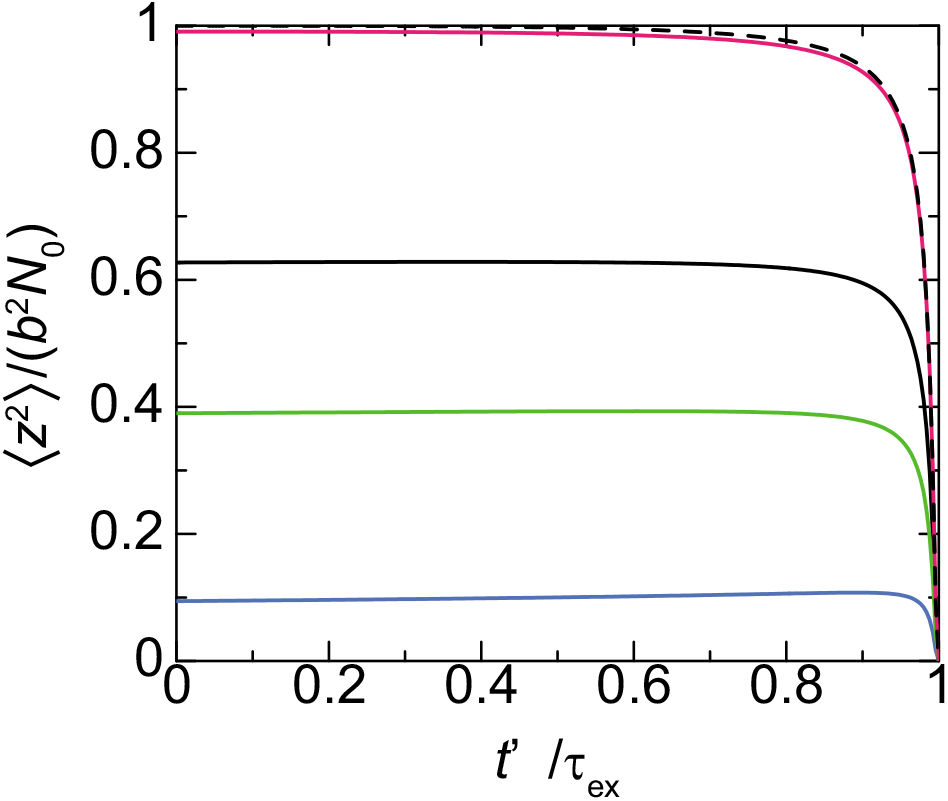
The mean square distance between the promoter and the surface is shown as a function of time *t′*/*τ*_ex_ elapsed since the loop extrusion process starts at *t*_0_/*τ*_*N*_0__ = 0.1 (cyan), 0.5 (light green), 1.0 (black), and 5.0 (magenta). We used *τ*_ex_/*τ*_*N*_0__ = 0.01. The black broken curve is calculated for an asymptotic limit, *t*_0_ → ∞.

**Figure 7:**
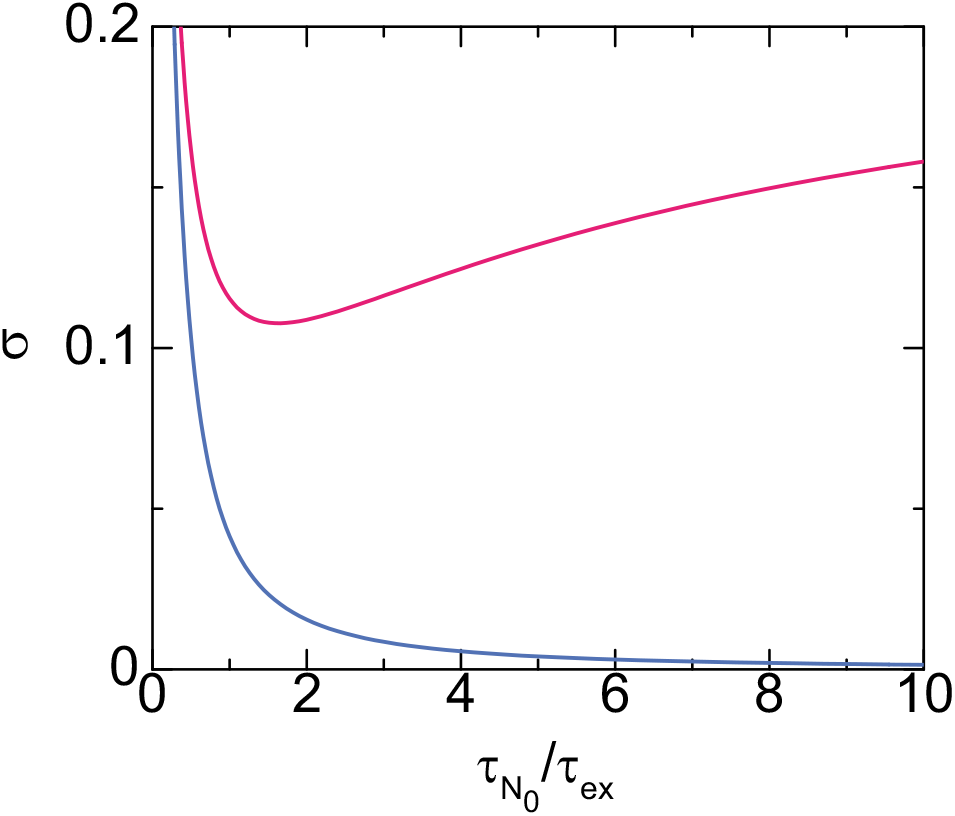
The contact probability of a promoter to the surface of the transcriptional condensate is shown as a function of the ratio *τ*_*N*_0__/*τ*_ex_ of time scales for cases in which the loop extrusion is active (magenta) and not active (cyan). We used *α*_elo_ = 0.3 and *ρ*/*K*_elo_ = 2.2 for the calculations. For the magenta line we set 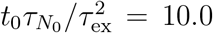, 2*k*_B_*Tτ_m_*/(*ζ*_0_*a*^2^) = 5.0 (the cyan line corresponds to 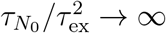). The ratio *τ*_*N*_0__/τ_ex_ scales linear to the number *N*_0_ of units in the linker chromatin, whereas the time scale 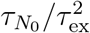 does not depend on the number *N*_0_ of units.

Our theory therefore predicts that the average loading time *τ*_on_ of cohesin and the relaxation time *τ*_*N*_0__ of the linker chromatin are the important time scales that determine the distribution of the position of the promoter.

### Contact probability of promoters to transcriptional condensate

The solution of eq. (14) for the steady state predicts that the factor 〈*e*^−*nϵ*/(*k*_B_*T*)^〉_*n*_ is derived as

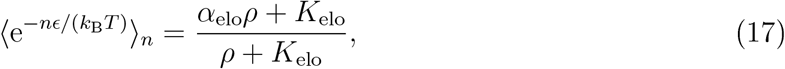

where *K*_elo_ (= λ_elo_*K*_ini_/(λ_elo_ + λ_ini_)) is the effective equilibrium constant. The factor *α*_elo_ has the form

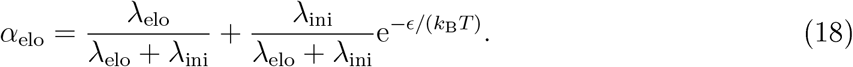

Eq. (17) has the asymptotic form 〈*e*^−*nϵ*/(*k*_B_*T*)^〉_*n*_ = e^−*ϵ*(*k*_B_*T*)^ for small rate constant λ_elo_ (because the promoter is occupied by pol II in the initiation state most of time) and 〈*e*^−*nϵ*/(*k*_B_*T*)^〉_*n*_ = 1 for large rate constant λ_elo_ (because the promoter is not occupied by pol II in the initiation state most of time).

In this section, we treat cases in which the concentration *ρ* of Pol II in the condensate is constant because the number of genes anchored to the condensate is small and the concentration *ρ* is relaxed to the equilibrium value in a relatively short time. For cases in which the loop extrusion is inhibited, the contact probability σ decreases monotonically as the number *N*_0_ of units in the linker chromatin increases, see the cyan line in fig. 8 (the ratio *τ*_*N*_0__/*τ*_ex_ of time scales is proportional to the number *N*_0_ of units). This is expected from the polymer physics: the linker chromatin acts an entropic spring which anchors the promoter to the surface of the transcription condensate and the stiffness of spring decreases as the number *N*_0_ of units in the linker chromatin increases.^29,30^ In sharp contrast, for cases in which the loop extrusion is active, the contact probability *σ* is a non-monotonic function of the number *N*_0_ of units in the linker chromatin, see the magenta line in fig. 8. In fact, the contact probability *σ* increases with the number *N*_0_ of units (for large enough *N*_0_). This is because the loop extrusion process pulls the promoter towards the surface of the condensate with a constant rate and the relaxation time *τ*_*N*_0__, with which the promoter stays in the proximity to the surface, increases as the number *N*_0_ of units in the linker chromatin increases, see eq. (16). On the other hand, the contact probability *σ* approaches asymptotically to the equilibrium values as the number *N*_0_ of segment decreases.

**Figure 8:**
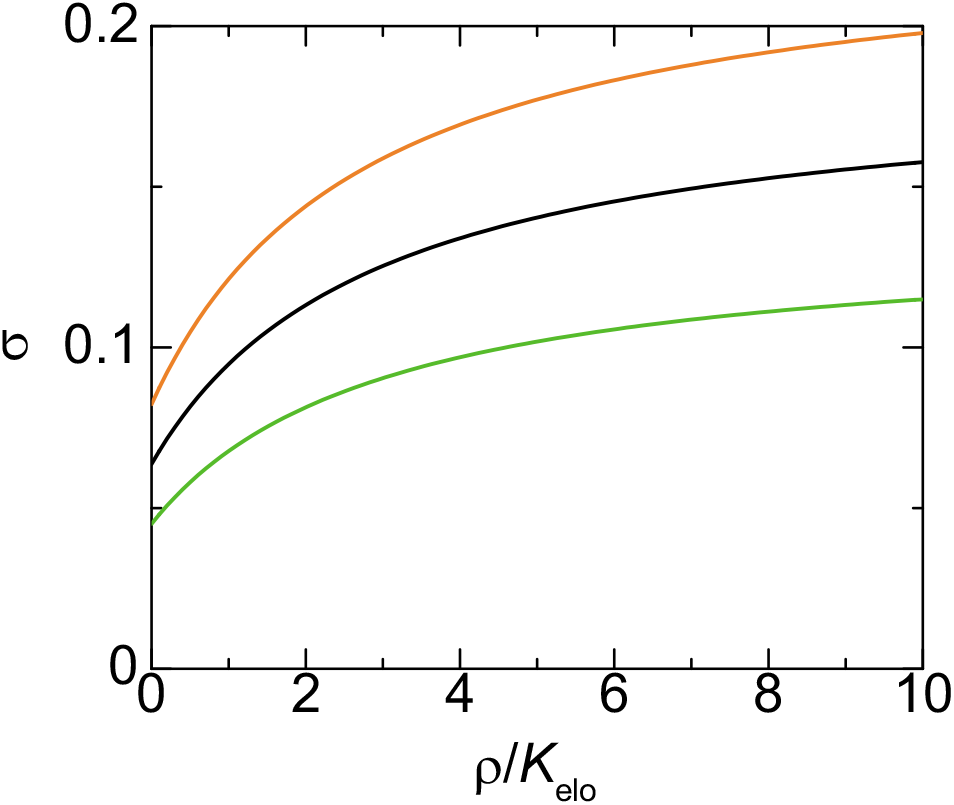
The contact probability of a promoter to the surface of the transcriptional condensate is shown as a function of the concentration *ρ* of Pol II in the condensate for *τ*_*N*_0__/*τ*_ex_ = 1.0 (light green), 10.0 (black), 20.0 (orange). We used *α*_elo_ = 0.3, 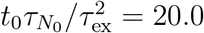, 2*k*_B_*Tτ_m_*/(*ζ*_0_*a*^2^) = 5.0 for the calculations.

The contact probability *σ* is represented as

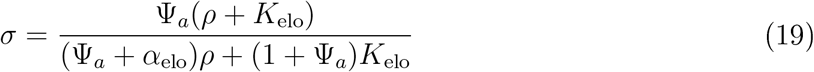

by using eq. (17), see fig. 8. The contact probability *σ* thus increases as the concentration *ρ* of Pol II increases (*α*_elo_ < 1 for ε > 0, see eq. (18), and the factor Ψ_a_ does not depend on the concentration *ρ* of Pol II). The contact probability has an asymptotic form *σ* = Ψ_*a*_/(Ψ_*a*_+*α*_elo_) for large concentrations *ρ*. The asymptotic probability reaches unity only for λ_elo_/λ_ini_ → 0 because Pol II tethers the promoter to the transcriptional condensate only in the initiation state. Our theory therefore predicts that the concentration of pol II in the condensate and the elongation rate are also important factors that determine the contact probability of the gene promoter to the transcriptional condensate.

## Discussion

Our theory predicts the contact probability of a promoter of a gene to the surface of a transcriptional condensate when the gene is anchored to the condensate via a superenhancer. This theory treats a relatively large transcriptional condensate, which is stable for the experimental time scale.^24^ Transcription machineries in the condensate are available to the promoter of the gene when the promoter is bound to the condensate. The contact probability therefore corresponds to the ratio of time in which the transcription of the gene is active during the transcription bursting. For cases in which the loop extrusion is inhibited, the contact probability decreases as the number of chromatin units in the linker chromatin increases. This reflects the fact that the linker chromatin acts as an entropic spring which anchors the promoter to the surface of the condensate and the stiffness of the spring decreases as the number of chromatin units in the linker chromatin increases. In contrast, for cases in which the loop extrusion is active, the contact probability of the promoter to the transcriptional condensate increases as the number of chromatin units in the linker chromatin increases as long as the promoter and the enhancer are in the same TAD. This is because the loop extrusion process drags the promoter to the surface with a constant rate and the relaxation time with which the promoter stays at the proximity to the surface increases as the number of chromatin units in the linker chromatin increases. This situation is very different from the static loops assumed in other theories.^8,9^

We used a couple of assumptions to simplify the theory: First, the binding and unbinding dynamics of the gene promoters is rate limited. It is motivated by the fact that among the genes that are at the proximity to a condensate and move together with the condensate, only 20 % of them colocalize with the condensate.^24^ If the binding of the gene promoters is diffusion limited, the unbinding of these promoters is a rare event: most of gene promoters would bind to the condensate in the steady state (or the transcriptional condensate observed in ref.^24^ is in the transient state). The limiting process may also depend on the transcription factors that bind to the gene promoter. Second, cohesin is trapped at the surface of the condensate. This is probably the case because the loop extrusion starts from the (super) enhancer^20^ and mediators may act as ‘boundary elements’ that stop the loop extrusion by cohesin.^11,12^ Single molecule experiments revealed that cohesin shows symmetric loop extrusion when there are no boundary elements.^18^ However, Hi-C experiments showed the signature of asymmetric loop extrusion, probably because cohesin loading sites are the proximity to boundary elements in such cases.^36^ Indeed, our main prediction reflects the dynamics of the linker chromatin in the relaxation process and is not very sensitive to the details of the dynamics in the loop extrusion process, as long as the average loading time *τ*_on_ is larger than the time scale *τ*_ex_ of the loop extrusion process. Third, the linker chromatin shows repulsive interactions with the transcriptional condensate. This treatment is motivated by the fact that chromatin tends to be excluded from the transcription condensate. ^25,26^ Fourth, there are at most one cohesin in the linker chromatin. The cases in which there are multiple cohesin molecules in the linker chromatin are treated by using *N*_p_ = *N*_0_*τ*_on_/*τ*_ex_ and *N*(*t*) = *N*_0_(*τ*_on_ − *t*)/*τ*_ex_. Fifth, we treated a relatively large condensate, which is stable for the experimental time scale (with which mouse embryonic stem cells are differentiated). ^24^ The lifetime of small transcriptional condensates is in the order of 10 s. The assembly and disassembly of such condensates may be coupled with processes involved in the transcription dynamics, such as the phospholylation of the C terminal domain of pol II^41^ and synthesis of RNA.^12^ We also assumed that the concentration of pol II in the condensate does not depend on the contact probability of gene promoters. Pol II with hyper-phospholylated C terminal domains does not have affinity to transcriptional condensates. ^41^ The pol II in the condensate is thus determined by the transcription rate and the relaxation time with which the concentration of pol II in the condensate relaxes to the equilibrium concentration. We assumed that the relaxation time with respect to the concentration of pol II is small, relative to the transcription rate, which is probably the cases in large transcriptional condensates. ^24^ Sixth, the contact probability of a promoter to the condensate is not affected by other gene promoters. It is of interest to theoretically predict 1) the coupling between the assembly/disassembly of transcriptional condensates and 2) the transcription dynamics and the interaction between gene promoters by using an extension of our theory.

There are growing number of researches on phase separation in biological systems. ^43^ Many researches emphasize the fact that the mutivalent interactions between intrinsically disordered domains of proteins play an important role in the formation of condensates ^43^ and that speciñc proteins and RNA are localized in condensates. ^24^ There is another important aspect of phase separation: phase separation creates interfaces. Superenhancers are anchored at the interface between a transcriptional condensate and the nucleoplasm. Our theory predicts that the promoters of the target genes of the superenhancers stay at the proximity to the interface because the loop extrusion process draggs the promoters to the interface and the slow dynamics of the linker chromatin between the promoter and the enhancer. Interfaces are asymmetric and (quasi-) 2d systems. Elucidating the roles played by interfaces in the biochemical reactions is an interesting avenue of phase separation researches in biological systems.

## Acknowledgement

This work was supported by JST, PRESTO Grant Number JPMJPR18KA (T.Y.), by JSPS KAKENHI Grant Number 18K03558 (T.Y.), by MEXT KAKENHI Grant Number JP20H05934 (T.Y.) and JP18H05529 (T.S.), and the Deutsche Forschungsgemeinschaft (DFG, German Research Foundation) under Germany’s Excellence Strategy - EXC 2068 - 390729961 - Cluster of Excellence Physics of Life of TU Dresden (H.S.). T.Y. acknowledges Tomoko Nishiyma (Nagoya University), Tsuyoshi Terakawa (Kyoto University), and Naomichi Takemata (Kyoto University) for fruitful discussion.

