## Supplementary Files for "Slow chromatin dynamics enhances promoter accessibility to transcriptional condensates"

### Supplementary File: Slow chromatin dynamics enhance promoter accessibility to transcriptional condensates

E-mail:

#### S1 Probability distribution function at (local) equilibrium

The probability distribution function $\psi_{\mathrm{eq}}(z)$ at equilibrium has the form

$$\psi_{\mathrm{eq}}(z) = \frac{q(z,N)q^*(z,N)}{\int_0^\infty dz q(z,N)q^*(z,N)}. \tag{S1}$$

The functions $q(z,s)$ and $q^*(z,s)$ are the solutions of the Edwards equation

$$\frac{\partial}{\partial s}q(z,s) = \frac{b^2}{6}\frac{\partial^2}{\partial z^2}q(z,s) \tag{S2}$$

$$\frac{\partial}{\partial s}q^*(z,s) = \frac{b^2}{6}\frac{\partial^2}{\partial z^2}q^*(z,s), \tag{S3}$$

where $z$ is the distance from the surface of the condensate and $s$ is the coordinate along the chromatin chain. $b$ denotes the Kuhn length. Eq. (S1) is the probability distribution function of the $N$-th unit in a chromatin chain, which is tethered to the surface of the condensate at $s = 0$ and $s = N_{\mathrm{p}}$. The boundary conditions for eqs. (S2) and (S3) are therefore

$$q(z,0) = q^*(z,N_{\mathrm{p}}) = \delta(z-\epsilon) \tag{S4}$$

$$q(0,s) = q^*(0,s) = 0. \tag{S5}$$

Eqs. (S2) - (S5) imply $q^*(z,s) = q(z, N_{\mathrm{p}} - s)$.

By separating the variables $q(z,s) = R(z)S(s)$, eq. (S2) is decomposed to equations for $R(z)$ and $S(s)$

$$\frac{\partial}{\partial s}S(s) = -\lambda S(s) \tag{S6}$$

$$\frac{\partial^2}{\partial z^2}R(z) = -\frac{6\lambda}{b^2}R(z). \tag{S7}$$

The solutions of eqs. (S6) and (S7) has the form

$$S(s) = \mathrm{e}^{-\lambda s} \tag{S8}$$

$$R(z) = \sin(kz) \tag{S9}$$

with

$$\lambda = \frac{k^2 b^2}{6}. \tag{S10}$$

We omitted $\cos(kz)$ from the solution of eq. (S9) by using the boundary condition, eq. (S5). The solution of eq. (S2) has the form

$$\begin{aligned} q(z,s) &= \frac{2}{L}\sum_k \sin(k\epsilon)\sin(kz)\mathrm{e}^{-k^2b^2s/6} \\ &= \frac{1}{\pi}\int_{-\infty}^{\infty} dk\ \sin(k\epsilon)\sin(kz)\mathrm{e}^{-k^2b^2s/6} \\ &= \sqrt{\frac{3}{2\pi b^2 s}}\left(\mathrm{e}^{-3(z-\epsilon)^2/(2b^2s)} - \mathrm{e}^{-3(z+\epsilon)^2/(2b^2s)}\right) \end{aligned} \tag{S11}$$

where we used eq. (S4) to derive the first line of eq. (S11). In the limit $\epsilon \to 0$, eq. (S11) has the asymptotic form

$$q(z,s) = \sqrt{\frac{3}{2\pi b^2 s}}\frac{6\epsilon}{b^2 s} z\mathrm{e}^{-3z^2/(2b^2s)}. \tag{S12}$$

In a similar manner, the function $q^*(z,s)$ is derived in the form

$$q^*(z,s) = \sqrt{\frac{3}{2\pi b^2(N_\mathrm{p}-s)}}\frac{6\epsilon}{b^2(N_\mathrm{p}-s)} z\mathrm{e}^{-3z^2/(2b^2(N_\mathrm{p}-s))}. \tag{S13}$$

The probability distribution function $\psi_\mathrm{eq}(z)$ at equilibrium has the form

$$\psi_\mathrm{eq}(z) = \frac{4}{\sqrt{\pi}}\frac{z^2}{(2l_\mathrm{p}^2)^{3/2}}\mathrm{e}^{-z^2/(2l_\mathrm{p}^2)} \tag{S14}$$

with

$$l_{\mathrm{p}}^2 = \frac{b^2}{3}\frac{N(N_{\mathrm{p}} - N)}{N_{\mathrm{p}}}. \tag{S15}$$

Eq. (S14) is reduced to the probability distribution function of the position $z$ of a promoter in a very long chromatin for $N_{\mathrm{p}} \to \infty$, see eq. (2).

With the loop extrusion process, the number $N_{\mathrm{p}}(t)$ of units in the loop, in which the chromatin section between the promoter and the enhancer is included, increases with time and has the form

$$N_{\mathrm{p}}(t) = N_0 \left(1 + \frac{t}{\tau_{\mathrm{ex}}}\right), \tag{S16}$$

where $N_0$ is the number of units in the linker chromatin between the promoter and the enhancer and $\tau_{\mathrm{ex}}$ is the time scale, with which the linker chromatin is extruded, see eq. (5) in the main article. The number $N(t)$ of units in the arm region is $N_0$ in the relaxation process, $0 < t < t_0$, and has the form

$$N(t') = N_0 \left(1 - \frac{t'}{\tau_{\mathrm{ex}}}\right), \tag{S17}$$

in the loop extrusion process, $t_0 < t < t_0 + \tau_{\mathrm{ex}}$, see eq. (6) in the main article. $t'$ $(= t - t_0)$ is the time elapsed since the loop extrusion starts. The probability distribution function at the local equilibrium is derived by substituting eqs. (S16) and (S17) into eq. (S15), see eqs. (7) and (8) in the main article.

#### S2 Probability distribution function in relaxation process

The time evolution of the probability distribution function $\psi(z,t)$ is given by the Smoluchowski equation

$$\frac{\partial}{\partial t}\psi(z,t) = \frac{k_{\mathrm{B}}T}{N\zeta}\frac{\partial}{\partial z}\left[\frac{\partial}{\partial z}\psi(z,t) + \frac{\partial}{\partial z}\left(\frac{U_{\mathrm{eff}}(z)}{k_{\mathrm{B}}T}\right)\psi(z,t)\right], \tag{S18}$$

where $U_{\mathrm{eff}}(z)$ is the effective potential that has the form

$$\frac{U_{\mathrm{eff}}(z)}{k_{\mathrm{B}}T} = \frac{z^2}{2l_{\mathrm{p}}^2} - 2\log z, \tag{S19}$$

see eqs. (1) - (4) in the main article. We derive the solution $\psi(z,t)$ of eq. (S18) in the form

$$\psi(z,t) = \frac{4}{\sqrt{\pi}}\frac{1}{(2l_{\mathrm{p}}^2)^{3/2}}\Phi(z,t)\mathrm{e}^{-U_{\mathrm{eff}}(z)/(k_{\mathrm{B}}T)}. \tag{S20}$$

Substituting eq. (S20) into eq. (S18) leads to the form

$$\frac{\partial}{\partial t}\Phi(z,t) = \frac{k_{\mathrm{B}}T}{N\zeta}\left[\frac{\partial^2}{\partial z^2}\Phi(z,t) - \left(\frac{z}{l_{\mathrm{p}}^2} - \frac{2}{z}\right)\frac{\partial}{\partial z}\Phi(z,t)\right]. \tag{S21}$$

By separating the variables $\Phi(z,t) = Z(z)T(t)$, eq. (S21) is decomposed to equations for $Z(z)$ and $T(t)$

$$\frac{\partial}{\partial t}T(t) = -\omega T(t) \tag{S22}$$

$$\frac{\partial^2}{\partial z^2}Z(z) - \left(\frac{z}{l_{\mathrm{p}}^2} - \frac{2}{z}\right)\frac{\partial}{\partial z}Z(z) = -\frac{N\zeta\omega}{k_{\mathrm{B}}T}Z(z). \tag{S23}$$

With the variable transformation

$$\xi = \frac{z^2}{2l_{\mathrm{p}}^2}, \tag{S24}$$

eq. (S23) is rewritten in the form

$$\xi Z''(\xi) + \left(\frac{3}{2} - \xi\right) Z'(\xi) + \frac{N\zeta l_{\mathrm{p}}^2 \omega}{2k_{\mathrm{B}}T} Z(\xi) = 0. \tag{S25}$$

Eq. (S25) is the Laguerre equation. The solution of eq. (S25) has the form

$$Z_n(\xi) = L_n^{1/2}(\xi) \tag{S26}$$

$$\omega_n = \frac{2k_{\mathrm{B}}T}{N\zeta l_{\mathrm{p}}^2} n, \tag{S27}$$

where $L_n^{1/2}(\xi)$ ($n = 0,\ 1,\ 2,\ \cdots$) is the Laguerre polynomials of the $n$-th rank. We rewrite $Z(\xi)$ and $\omega$ to $Z_n(\xi)$ and $\omega_n$, respectively, because these functions depend on the index $n$. The solution of eq. (S22) is readily derived as

$$T_n(t) = T_n(0)\mathrm{e}^{-\omega_n t}. \tag{S28}$$

We rewrite $T(t)$ to $T_n(t)$ because this function depends on the index $n$. The solution of eq. (S21) therefore has the form

$$\Phi(z,t) = \sum_{n=0}^{\infty} C_n(0) L_n^{1/2}(\xi) \mathrm{e}^{-\omega_n t}, \tag{S29}$$

where $C_n(0)$ ($n = 0,\ 1,\ 2,\ \cdots$) are constants that are determined by the initial condition. By substituting eq. (S29) into eq. (S20), the general solution of the probability distribution function $\psi(z,t)$ is derived as

$$\psi(z,t)dz = \sum_{n=0}^{\infty} C_n(0) L_n^{1/2}(\xi) \xi^{1/2} \mathrm{e}^{-\xi} \mathrm{e}^{-\omega_n t} d\xi. \tag{S30}$$

By using the probability distribution function $\psi_0(z)$ ($\equiv \psi(z,0)$) at $t = 0$, the coefficients

$C_n(0)$ are written as

$$C_n(0) = \frac{\Gamma(n+1)}{\Gamma(n+3/2)} \int_0^\infty dz\, L_n^{1/2}(\xi)\psi_0(z). \tag{S31}$$

Eq. (S31) is derived by using the orthogonality of the Laguerre polynomials

$$\int_0^\infty d\xi\, \mathrm{e}^{-\xi}\xi^{1/2} L_n^{1/2}(\xi) L_m^{1/2}(\xi) = \begin{cases} 0 & (m \neq n) \\ \frac{\Gamma(n+3/2)}{\Gamma(n+1)} & (m = n). \end{cases} \tag{S32}$$

We first treat a simple case in which the promoter is located at $z = 0$ at $t = 0$, $\psi_0(z) = \delta(z)$. In this case, the coefficients $C_n(0)$ are given by

$$C_n(0) = \frac{2}{\sqrt{\pi}}, \tag{S33}$$

where we used the relationship

$$L_n^{1/2}(0) = \frac{2}{\sqrt{\pi}} \frac{\Gamma(n+3/2)}{\Gamma(n+1)}. \tag{S34}$$

Substituting eq. (S33) into eq. (S30) leads to the form

$$\begin{aligned} \psi(z,t)dz &= \frac{2}{\sqrt{\pi}} \sum_{n=0}^{\infty} L_n^{1/2}(\xi)\xi^{1/2}\mathrm{e}^{-\xi}\mathrm{e}^{-\omega_n t} d\xi \\ &= \frac{2}{\sqrt{\pi}} \frac{\xi^{1/2} d\xi}{(1-\mathrm{e}^{-t/\tau_\mathrm{p}})^{3/2}} \mathrm{e}^{-\xi/(1-\mathrm{e}^{-t/\tau_\mathrm{p}})} \\ &= \frac{4}{\sqrt{\pi}} \frac{z^2 dz}{(2l_\mathrm{p}^2 r_\mathrm{s}(t))^{3/2}} \mathrm{e}^{-z^2/(2l_\mathrm{p}^2 r_\mathrm{s}(t))}, \end{aligned} \tag{S35}$$

with

$$r_\mathrm{s}(t) = 1 - \mathrm{e}^{-t/\tau_\mathrm{p}}. \tag{S36}$$

The relaxation time $\tau_{\rm p}$ has the form

$$\tau_{\rm p} = \frac{N\zeta l_{\rm p}^2}{2k_{\rm B}T}. \tag{S37}$$

We used $\omega_n = n/\tau_{\rm p}$ and the generating function of the Laguerre polynomials

$$\sum_{n=0}^{\infty} L_n^{1/2}(\xi)x^n = \frac{1}{(1-x)^{3/2}}{\rm e}^{-\xi x/(1-x)} \tag{S38}$$

to derive the second line of eq. (S35).

With loop extrusion present, the time evolution equation of the relaxation process is given by

$$\begin{aligned} & \frac{\partial}{\partial t}\psi(z,t;N,N_{\rm p}) + \frac{1}{\tau_{\rm m}}\frac{\partial}{\partial N_{\rm p}}\psi(z,t;N,N_{\rm p}) \\ = & \frac{k_{\rm B}T}{N\zeta}\frac{\partial}{\partial z}\left[\frac{\partial}{\partial z}\psi(z,t;N,N_{\rm p}) + \frac{\partial}{\partial z}\left(\frac{U_{\rm eff}(z,t)}{k_{\rm B}T}\right)\psi(z,t;N,N_{\rm p})\right], \end{aligned} \tag{S39}$$

see eq. (10) in the main article. By using the characteristic curves

$$\frac{dN(t)}{dt} = 0 \tag{S40}$$

$$\frac{dN_{\rm p}(t)}{dt} = \frac{1}{\tau_{\rm m}}, \tag{S41}$$

eq. (S39) is rewritten as

$$\frac{\partial}{\partial t}\psi(z,t;N,N_{\rm p}(t)) = \frac{k_{\rm B}T}{N\zeta}\frac{\partial}{\partial z}\left[\frac{\partial}{\partial z}\psi(z,t;N,N_{\rm p}(t)) + \frac{\partial}{\partial z}\left(\frac{U_{\rm eff}(z,t)}{k_{\rm B}T}\right)\psi(z,t;N,N_{\rm p}(t))\right]. \tag{S42}$$

The solutions of eqs. (S40) and (S41) are $N = N_0$ and eq. (S16).

We derive the solution of eq. (S42) in the form

$$\psi(z,t) = \frac{4}{\sqrt{\pi}}\frac{z^2}{(2l_{\mathrm{eq}}^2 r_{\mathrm{p}}(t))^{3/2}}\mathrm{e}^{-z^2/(2l_{\mathrm{eq}}^2 r_{\mathrm{p}}(t))}. \tag{S43}$$

Substituting eq. (S43) into eq. (S42) leads to the form

$$\text{(Left side)} = \left(\frac{d}{dt}r_{\mathrm{p}}(t)\right)\left[-\frac{3}{2}\frac{1}{r_{\mathrm{p}}}+\frac{1}{2}\frac{z^2}{l_{\mathrm{eq}}^2 r_{\mathrm{p}}^2}\right]\psi(z,t) \tag{S44}$$

$$\text{(Right side)} = -\frac{2}{l_{\mathrm{eq}}^2}\frac{k_{\mathrm{B}}T}{N\zeta}\left(\frac{l_{\mathrm{eq}}^2}{l_{\mathrm{p}}^2}r_{\mathrm{p}}-1\right)\left[-\frac{3}{2}\frac{1}{r_{\mathrm{p}}}+\frac{1}{2}\frac{z^2}{l_{\mathrm{eq}}^2 r_{\mathrm{p}}^2}\right]\psi(z,t). \tag{S45}$$

Eq. (S42) is thus reduced to

$$\frac{d}{dt}r_{\mathrm{p}}(t) = -\frac{2}{l_{\mathrm{eq}}^2}\frac{k_{\mathrm{B}}T}{N\zeta}\left(\frac{l_{\mathrm{eq}}^2}{l_{\mathrm{p}}^2}r_{\mathrm{p}}(t)-1\right). \tag{S46}$$

The solution of eq. (S46) has the form

$$r_{\mathrm{p}}(t) = \frac{2k_{\mathrm{B}}T}{N_0\zeta l_{\mathrm{eq}}^2}\int_0^t dt_2\,\mathrm{e}^{-\int_{t_2}^t dt_1\,\frac{2k_{\mathrm{B}}T}{N_0\zeta l_{\mathrm{p}}^2(t_1)}}, \tag{S47}$$

where we used $r_{\mathrm{p}}(0) = 0$ for the initial condition (corresponding to $\psi(z,0) = \delta(z)$).

### S3 Probability distribution function in loop extrusion process

In the loop extrusion process, the time evolution equation of the probability distribution function has the form

$$\begin{aligned}&\frac{\partial}{\partial t}\psi(z,t;N,N_{\mathrm{p}}) - \frac{1}{\tau_{\mathrm{m}}}\frac{\partial}{\partial N}\psi(z,t;N,N_{\mathrm{p}}) + \frac{1}{\tau_{\mathrm{m}}}\frac{\partial}{\partial N_{\mathrm{p}}}\psi(z,t;N,N_{\mathrm{p}})\\ &= \frac{k_{\mathrm{B}}T}{N_0\zeta}\frac{\partial}{\partial z}\left[\frac{\partial}{\partial z}\psi(z,t;N,N_{\mathrm{p}}) + \frac{\partial}{\partial z}\left(\frac{U_{\mathrm{eff}}(z,t)}{k_{\mathrm{B}}T}\right)\psi(z,t;N,N_{\mathrm{p}})\right].\end{aligned} \tag{S48}$$

By using the characteristic curves

$$\frac{dN_{\rm p}(t)}{dt} = \frac{1}{\tau_{\rm m}} \tag{S49}$$

$$\frac{dN(t)}{dt} = -\frac{1}{\tau_{\rm m}}, \tag{S50}$$

eq. (S48) is rewritten in the form

$$\frac{\partial}{\partial t}\psi(z,t;N(t),N_{\rm p}(t)) = \frac{k_{\rm B}T}{N_0\zeta}\frac{\partial}{\partial z}\left[\frac{\partial}{\partial z}\psi(z,t;N(t),N_{\rm p}(t)) + \frac{\partial}{\partial z}\left(\frac{U_{\rm eff}(z,t)}{k_{\rm B}T}\right)\psi(z,t;N(t),N_{\rm p}(t))\right]. \tag{S51}$$

The solutions of eqs. (S49) and (S50) are eqs. (S16) and (S17), respectively.

We derive the solution of eq. (S51) in the form

$$\psi(z,t) = \frac{4}{\sqrt{\pi}}\frac{z^2}{(2l_{\rm eq}^2 r_{\rm ex}(t'))^{3/2}}{\rm e}^{-z^2/(2l_{\rm eq}^2 r_{\rm ex}(t'))}, \tag{S52}$$

where $t'$ ($= t - t_0$) is the time elapsed after the loop extrusion starts, see also eq. (S35). Substituting eq. (S52) into eq. (S51) leads to the form

$$(\text{Left side}) = \left(\frac{d}{dt}r_{\rm ex}\right)\left[-\frac{3}{2}\frac{1}{r_{\rm ex}} + \frac{1}{2}\frac{z^2}{l_{\rm eq}^2 r_{\rm ex}^2}\right]\psi(z,t) \tag{S53}$$

$$(\text{Right side}) = -\frac{2}{l_{\rm eq}^2}\frac{k_{\rm B}T}{N\zeta}\left(\frac{l_{\rm eq}^2}{l_{\rm p}^2}r_{\rm ex} - 1\right)\left[-\frac{3}{2}\frac{1}{r_{\rm ex}} + \frac{1}{2}\frac{z^2}{l_{\rm eq}^2 r_{\rm ex}^2}\right]\psi(z,t). \tag{S54}$$

Eq. (S52) is finally reduced to

$$\frac{d}{dt}r_{\rm ex}(t) = -\frac{2}{l_{\rm eq}^2}\frac{k_{\rm B}T}{N\zeta}\left(\frac{l_{\rm eq}^2}{l_{\rm p}^2(t)}r_{\rm ex}(t) - 1\right). \tag{S55}$$

The solution of eq. (S55) has the form

$$r_{\rm ex}(t') = r_{\rm p}(t_0){\rm e}^{-\int_0^{t'}dt_1\,\frac{2k_{\rm B}T}{N(t_1)\zeta l_{\rm p}^2(t_1)}} + \int_0^{t'}dt_2\frac{2k_{\rm B}T}{N(t_2)\zeta l_{\rm eq}^2}{\rm e}^{\int_{t'}^{t_2}dt_1\,\frac{2k_{\rm B}T}{N(t_1)\zeta l_{\rm p}^2(t_1)}}, \tag{S56}$$

where we set $r_{\rm ex}(t' = 0) = r_{\rm p}(t_0)$ for the initial condition.
